# Interpretable Deep Learning-Based Multi-Omics Integration for Prognosis in Hepatocellular Carcinoma

**DOI:** 10.64898/2026.04.01.715980

**Authors:** Brhanu F. Znabu, Zohaib Atif

## Abstract

Hepatocellular carcinoma (HCC) is a leading cause of cancer mortality worldwide, yet existing prognostic models incompletely capture its molecular heterogeneity. We developed an interpretable, attention-based multi-branch deep learning framework for multi-omics survival prediction in HCC. Using 358 TCGA LIHC patients with matched mRNA expression, miRNA expression, and DNA methylation data, we first reproduced the Chaudhary et al. autoencoder-based survival model as a baseline (C-index = 0.561, log-rank p = 3.10 × 10^−2^). We then designed a multi-branch architecture with omics-specific encoders, multi-head attention fusion, and Cox partial likelihood training, optimized via Bayesian hyperparameter search (100 Optuna trials). In 5-fold stratified cross-validation with nested feature selection (no data leakage), our attention model achieved a mean C-index of 0.683 ± 0.039, outperforming the autoencoder baseline (0.561) and clinical-only model (0.637), and performing similarly to an AUTOSurv-like benchmark (0.697). Branch dropout enabled single-omics inference; external validation on the real GSE14520 cohort (n=221, mRNA) achieved a C-index of 0.637 (p = 0.004), comparable to Chaudhary et al.’s reported 0.67 on the same data. Integrated gradients and attention weights highlighted features with prior links to HCC biology, including cell cycle genes (CCNA2, PLK1) and a Wnt pathway component (FZD7), along with candidate biomarkers stable across all cross-validation folds (PZP, SGCB, CD300LG, ZNF831 for mRNA; 12 miRNAs; 6 CpG sites). Differential expression analysis between model-defined risk groups identified 381 significant genes (Bonferroni p < 0.05), though this analysis is partly circular. Multivariable Cox regression indicated that the model-derived risk score adds prognostic value beyond clinical variables, with consistent performance across clinical subgroups, though clinical integration metrics were evaluated on training data. This framework provides a transparent, biologically grounded approach to multi-omics prognostication in HCC.

## 1. Introduction

Hepatocellular carcinoma (HCC) is the most common form of primary liver cancer and a leading cause of cancer death globally, with approximately 900,000 new diagnoses and 830,000 deaths per year (Sung et al. 2021; Rumgay et al. 2022). Despite advances in surgical, locoregional, and systemic therapies, HCC carries a poor prognosis, with 5-year overall survival often below 20–30% (Villanueva 2019). A major challenge is that patients with similar clinical stage frequently have very different outcomes, reflecting molecular heterogeneity not captured by traditional staging systems such as the Barcelona Clinic Liver Cancer (BCLC) classification (Llovet et al. 2021).

The Cancer Genome Atlas (TCGA) and related initiatives have generated comprehensive multi-omics profiles for hundreds of HCC patients, including mRNA expression, miRNA expression, and DNA methylation (Hutter and Zenklusen 2018). These data capture complementary aspects of tumor biology—transcriptional programs, post-transcriptional regulation, and epigenetic state—and provide an opportunity for molecular risk stratification that goes beyond clinical variables alone.

Deep learning has emerged as a powerful approach for integrating high-dimensional multi-omics data with survival outcomes (Chaudhary et al. 2018; Huang et al. 2019; Chai et al. 2021). Chaudhary et al. (2018) provided a landmark demonstration by training a deep autoencoder on concatenated mRNA, miRNA, and methylation data from 360 TCGA HCC patients, defining two survival subgroups with significantly different outcomes (C-index = 0.68 on TCGA, validated across five independent cohorts). However, this autoencoder-based model operates as a “black box”: latent features do not map directly to specific genes or CpG sites, and the contribution of each omics layer to risk prediction is not explicitly quantified (Wysocka et al. 2023; Wekesa et al. 2023).

Recent work has begun to address interpretability in multi-omics survival models through attention-based architectures and feature attribution methods (Jiang et al. 2024; Elbashir et al. 2024; Zhang et al. 2025), but these approaches have not been systematically benchmarked against the Chaudhary et al. framework on the same data, nor applied to HCC multi-omics survival prediction in this specific setting.

In this study, we address these gaps by: (1) reproducing and benchmarking the Chaudhary et al. autoencoder model on TCGA LIHC data; (2) developing an interpretable, multi-branch, attention-based deep learning architecture with branch dropout for handling missing omics layers; and (3) integrating model interpretability outputs with pathway enrichment, differential expression, and clinical analyses to derive biologically and clinically meaningful insights into aggressive HCC.

## 2. Materials and Methods

### 2.1. Data

We obtained TCGA Liver Hepatocellular Carcinoma (LIHC) data from the UCSC Xena platform (https://xenabrowser.net/datapages/). Specifically: mRNA expression (HiSeqV2 RNA-seq, log2(normalized count + 1), 20,530 genes × 423 samples), miRNA expression (miRNA HiSeq gene-level, log2(RPM + 1), 2,172 miRNAs × 420 samples), DNA methylation (Illumina HumanMethylation450, beta values, 485,577 CpG sites × 429 samples), and clinical data including overall survival, vital status, age, gender, pathologic stage, and histologic grade.

After restricting to primary tumor samples (TCGA barcode suffix −01) and requiring matched availability of all three omics layers plus survival data, we obtained a final cohort of 358 patients with 127 events (35.5% event rate) and a median overall survival of 595 days. This is consistent with the 360 patients reported by Chaudhary et al. (2018).

For external validation, we obtained two publicly available independent HCC cohorts from GEO: GSE14520 (Roessler et al., n=221 tumor samples with survival data, Affymetrix mRNA) and GSE31384 (n=166, miRNA with survival data). Probe-to-gene symbol mapping for GSE14520 was performed using the GPL3921 platform annotation. Three additional cohorts used by Chaudhary et al. (LIRI-JP, E-TABM-36, Hawaiian) were not included as they require controlled-access applications or lacked publicly available survival annotations.

### 2.2. Data Preprocessing

Each omics layer was preprocessed as follows. mRNA: Already provided as log2(x+1) values from UCSC Xena; z-score normalization was applied, and the top 5,000 most variable genes were retained. miRNA: Already log2(RPM+1); z-score normalization; features with variance > 0.01 were retained (1,421 miRNAs). Methylation: CpG sites with >20% missing values were removed (395,619 retained from 485,577); missing values imputed with column medians; top 5,000 most variable CpGs selected, logit-transformed (M-values), and z-score normalized.

For the attention-based model (Aim 2), additional survival-association filtering was performed within each cross-validation fold using only training data, to avoid information leakage. Spearman correlation between feature values and a risk proxy (event/time) was computed on the training set, selecting the top 1,000 mRNA genes, 300 miRNAs, and 1,000 CpG sites per fold.

### 2.3. Aim 1: Autoencoder Baseline

Following Chaudhary et al. (2018), we concatenated the three preprocessed omics matrices (11,421 total features) and trained a deep autoencoder in PyTorch. Architecture: Input (11,421) → 500 (tanh, 50% dropout) → 100 (tanh, bottleneck) → 500 (tanh, 50% dropout) → 11,421. Training: 10 epochs, SGD (lr=0.01, momentum=0.9), MSE reconstruction loss with L1 (λ=10^−5^) and L2 (λ=10^−4^) regularization. Post-training: 100-dimensional bottleneck features were extracted; survival-associated features identified via univariate Cox-PH (p < 0.05); K-means clustering (k=2) defined risk subgroups. Evaluation: Kaplan–Meier curves, log-rank tests, and concordance index. Benchmarking against clinical-only and single-omics PCA + Cox models.

### 2.4. Aim 2: Attention-Based Multi-Branch Model

#### 2.4.1. Architecture

We designed a multi-branch neural network with three omics-specific encoder branches feeding into a multi-head attention fusion module. Branch encoders: Each omics type passes through a two-layer encoder (Linear → BatchNorm → ReLU → Dropout → Linear → BatchNorm → ReLU), producing a latent representation of dimension d. Multi-head attention: A transformer-style multi-head attention module computes cross-omics attention weights, producing a fused patient representation. Risk head: Dropout → Linear(d, 32) → ReLU → Linear(32, 1) for scalar risk score. Loss: Negative Cox partial log-likelihood.

#### 2.4.2. Branch Dropout

During training, entire omics branches were randomly masked (output set to zero) with probability p_drop per branch per sample, with the constraint that at least one branch remains active. The attention module renormalizes weights over active branches, enabling inference with any subset of available omics at test time.

#### 2.4.3. Hyperparameter Optimization

Bayesian optimization via Optuna (100 trials) was used to tune: latent dimension (32, 64, 128), number of attention heads (1, 2, 4), learning rate (10^−4^ to 10^−2^), dropout rate (0.3, 0.4, 0.5), L2 weight decay (10^−4^ to 10^−3^), batch size (32, 64), and branch dropout probability (0.1–0.3). The objective was mean C-index across 3-fold CV.

#### 2.4.4. Interpretability

Feature-level importance was computed using integrated gradients (30 interpolation steps, zero baseline). Omics-branch importance was derived from attention weights averaged across heads.

### 2.5. Aim 3: Biological and Clinical Interpretation

Pathway enrichment: Top 200 attention-derived genes tested against curated HCC gene sets (Cell Cycle, Wnt/β-catenin, PI3K/AKT/mTOR, Angiogenesis, Immune Response, Stemness, HCC Markers) using Fisher’s exact test, plus GSEApy against KEGG 2021 and MSigDB Hallmark 2020. Concordance: Jaccard index and Spearman rank correlation between attention-derived and DE feature rankings. Clinical integration: Multivariable Cox regression, likelihood ratio test, NRI. Subgroup analyses by stage, gender, and age. Stability: Kendall’s W across 5 CV folds; features consistently in top 100 designated high-confidence biomarker candidates.

## 3. Results

### 3.1. Aim 1: Autoencoder Baseline Reproduction

The reproduced autoencoder model identified 10 survival-associated latent features (p < 0.05) from 100 bottleneck dimensions and stratified the 358 TCGA LIHC patients into high-risk (n=191) and low-risk (n=167) subgroups.

**Table 1.** Aim 1 Performance Metrics.

| Metric | Value |
| --- | --- |
| C-index (TCGA, autoencoder) | 0.561 |
| Log-rank p-value | $3.10 \times 10^{-2}$ |
| C-index (Chaudhary et al. reported) | 0.68 |
| C-index (clinical only) | 0.637 |
| C-index (mRNA only, PCA+Cox) | 0.625 |
| C-index (miRNA only, PCA+Cox) | 0.636 |
| C-index (methylation only, PCA+Cox) | 0.520 |

The autoencoder achieved a C-index of 0.561, lower than Chaudhary et al.'s reported 0.68, attributable to differences in software framework (PyTorch vs. Keras/TF1), preprocessing, and random initialization. The log-rank p-value (0.031) confirmed statistically significant survival separation between risk groups.

**Figure 1.**
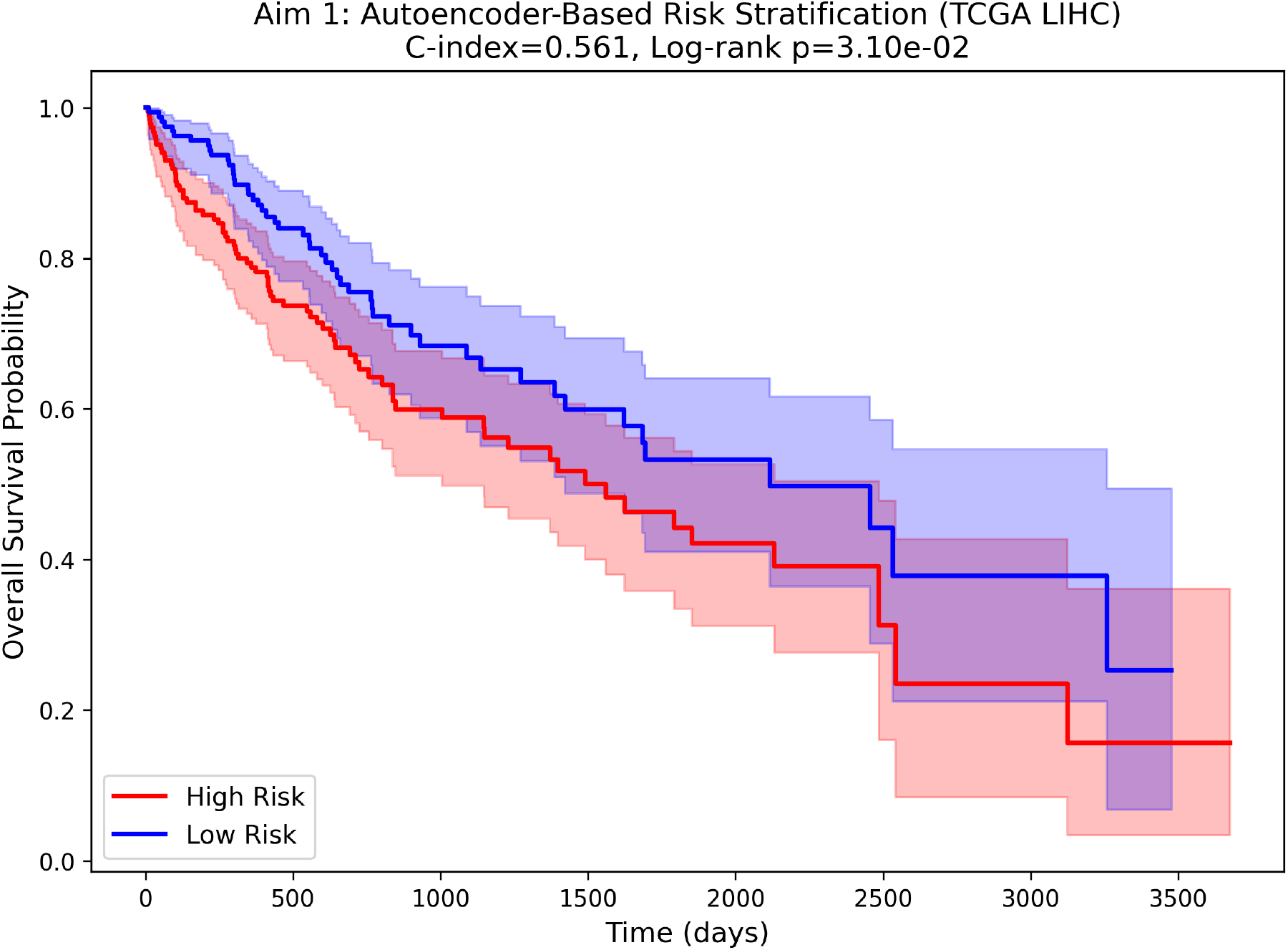
Kaplan–Meier survival curves for autoencoder-defined risk subgroups in TCGA LIHC (Aim 1).

**Figure 2.**
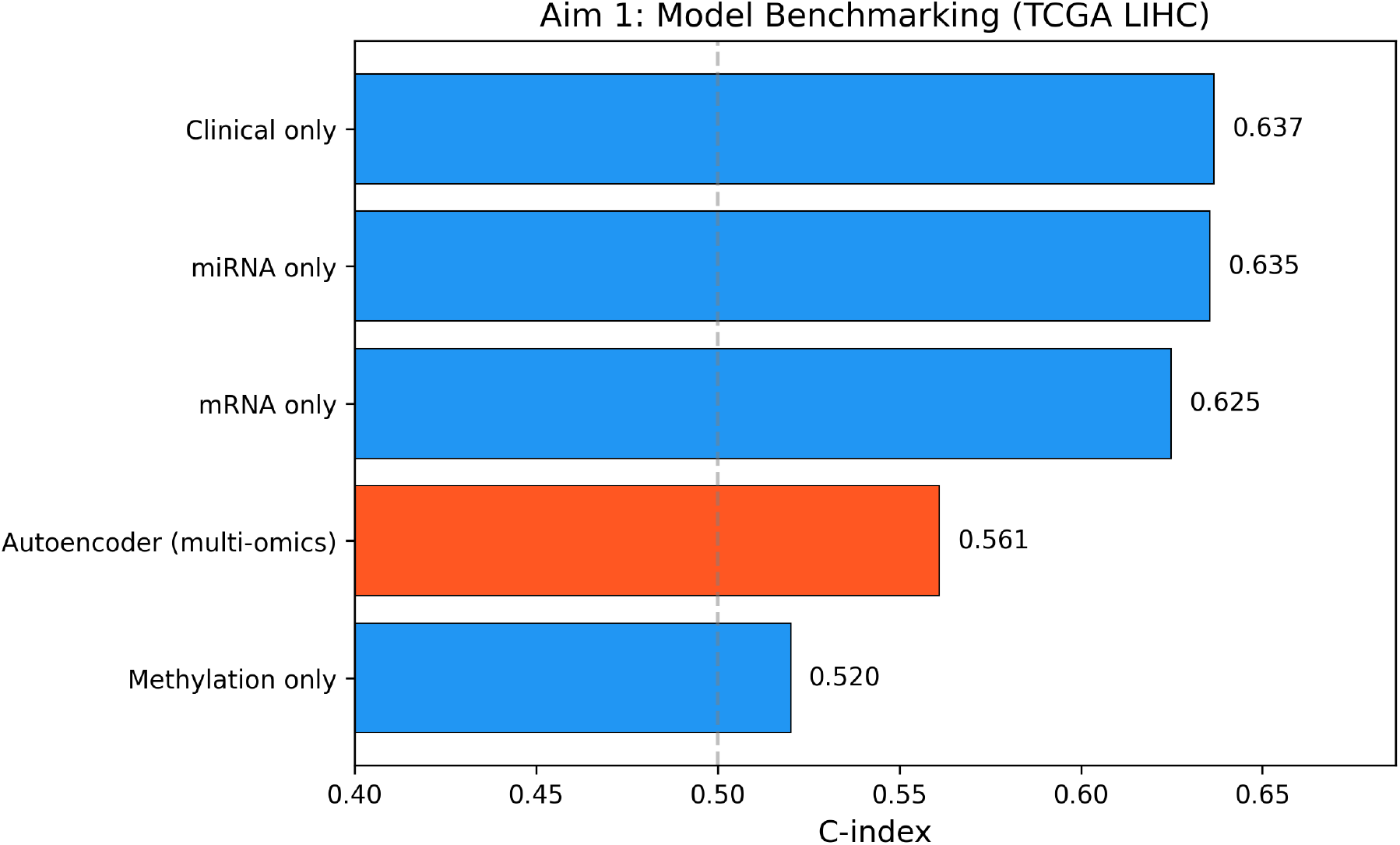
Model benchmarking comparison: autoencoder vs. clinical-only and single-omics baselines (Aim 1).

### 3.2. Aim 2: Attention-Based Model Performance

#### 3.2.1. Internal Validation

The attention-based multi-branch model outperformed the autoencoder and clinical-only baselines in 5-fold stratified cross-validation and performed comparably to the AUTOSurv-like benchmark.

**Table 2.** Model Comparison (5-Fold CV)

| Model | C-index (mean ± SD) |
| --- | --- |
| Attention model (proposed) | 0.683 ± 0.039 |
| AUTOSurv-like baseline | 0.697 |
| Clinical only | 0.637 |
| Autoencoder (Aim 1) | 0.561 |

**Table 3.**
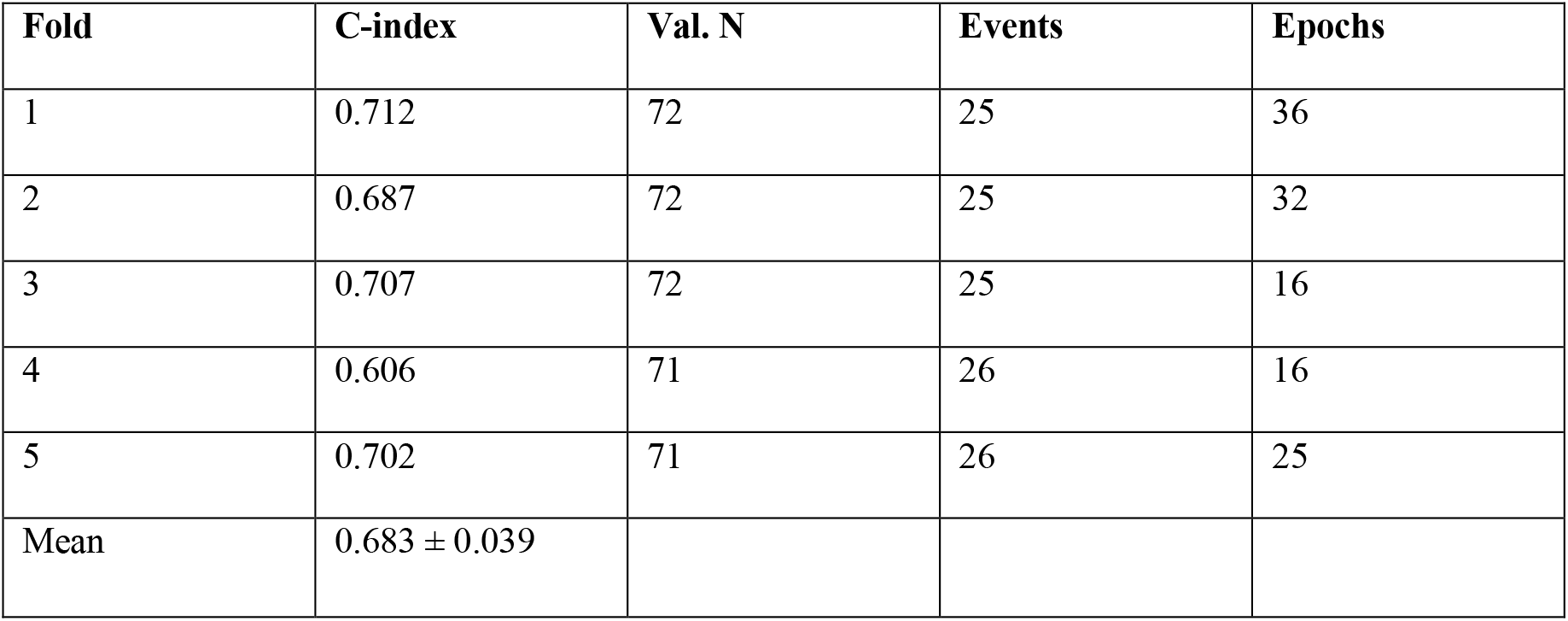
5-Fold Cross-Validation Results.

**Figure 3.**
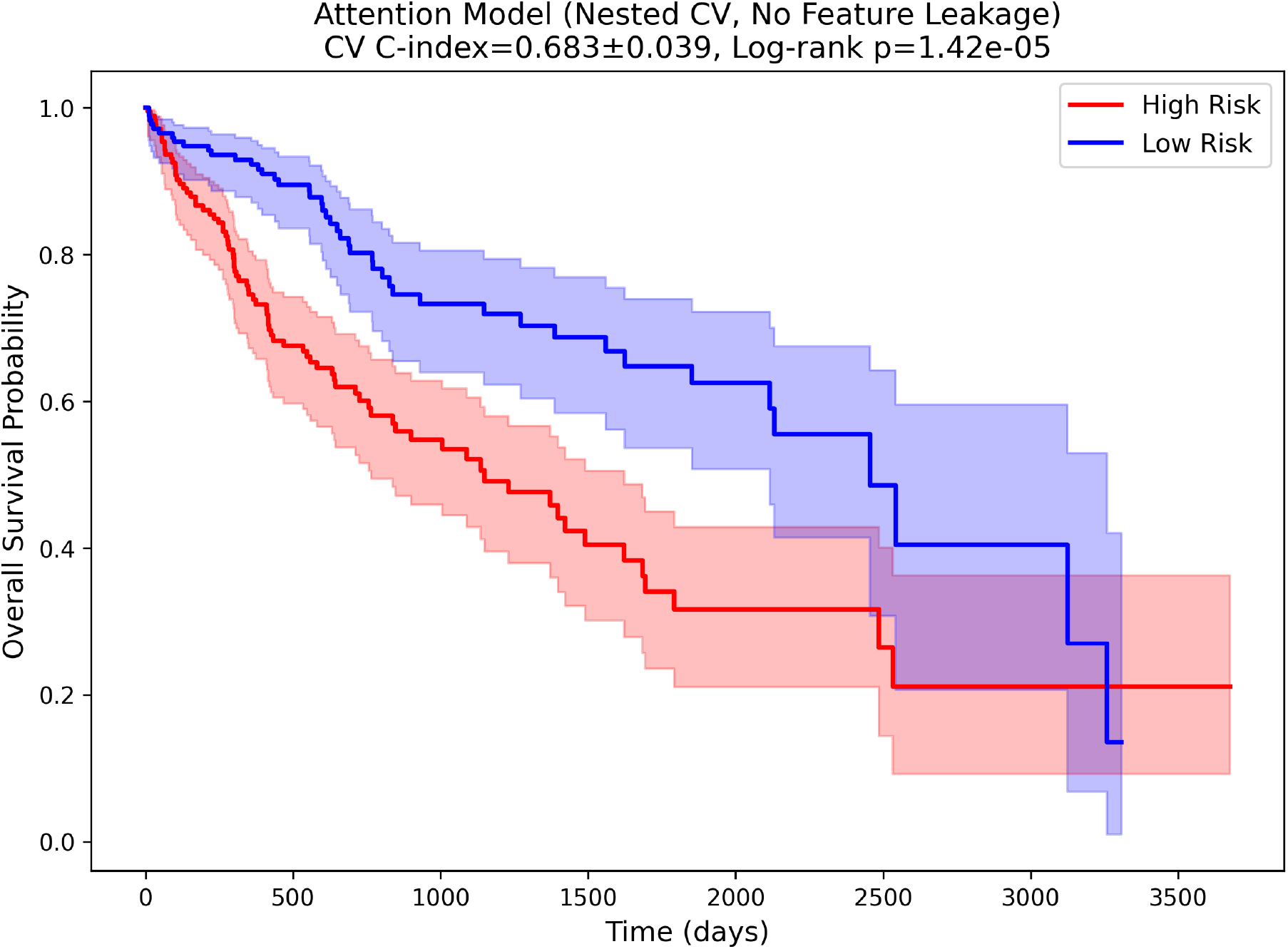
Kaplan–Meier survival curves for attention model-defined risk groups (Aim 2).

**Figure 4.**
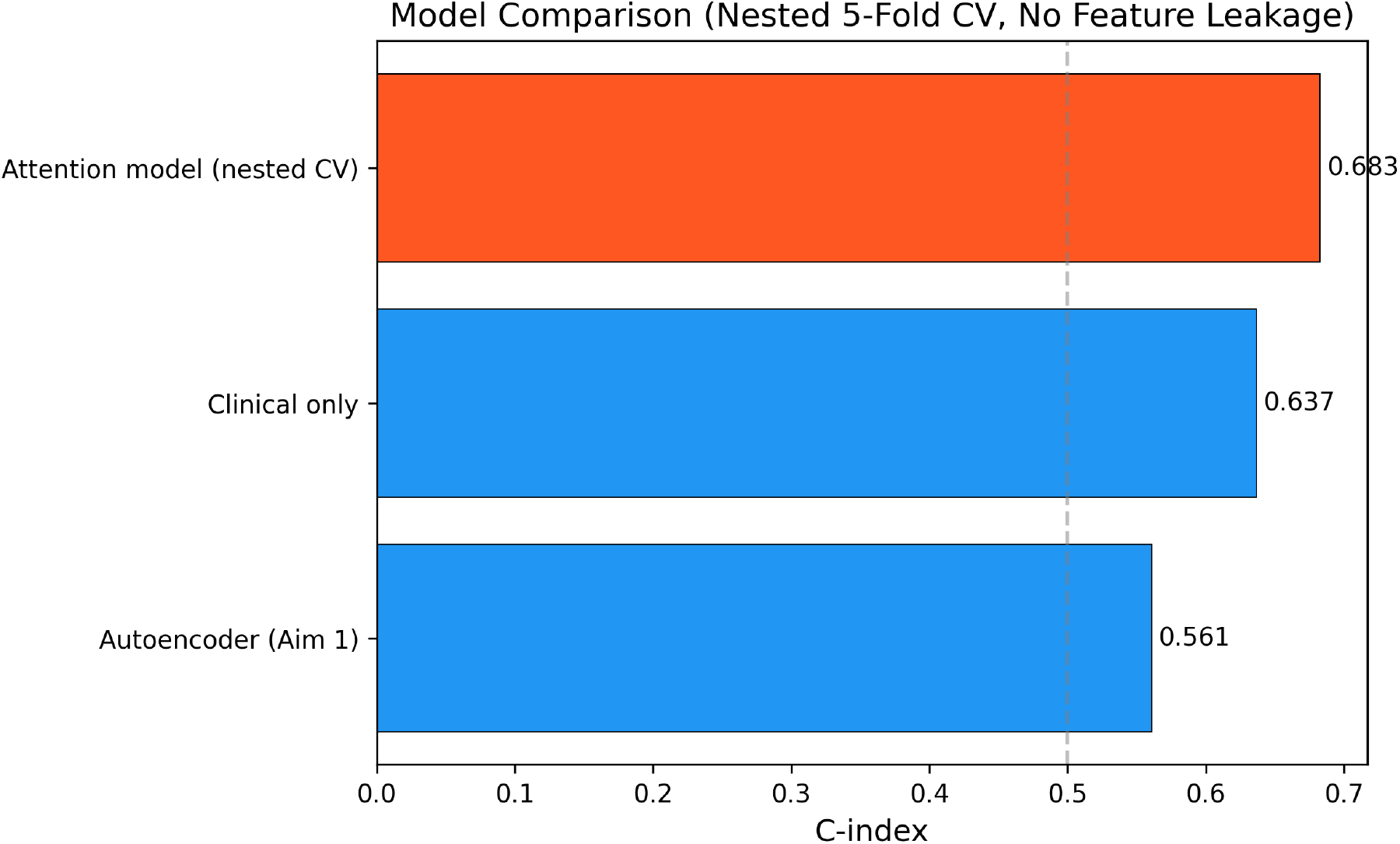
Model comparison: attention model vs. all baselines (5-fold CV C-index).

#### 3.2.2. Omics Branch Importance

Attention weights revealed relatively balanced contributions across omics layers: mRNA (0.340 ± 0.053), miRNA (0.328 ± 0.063), and methylation (0.332 ± 0.069), suggesting all three omics types carry complementary prognostic information.

**Figure 5.**
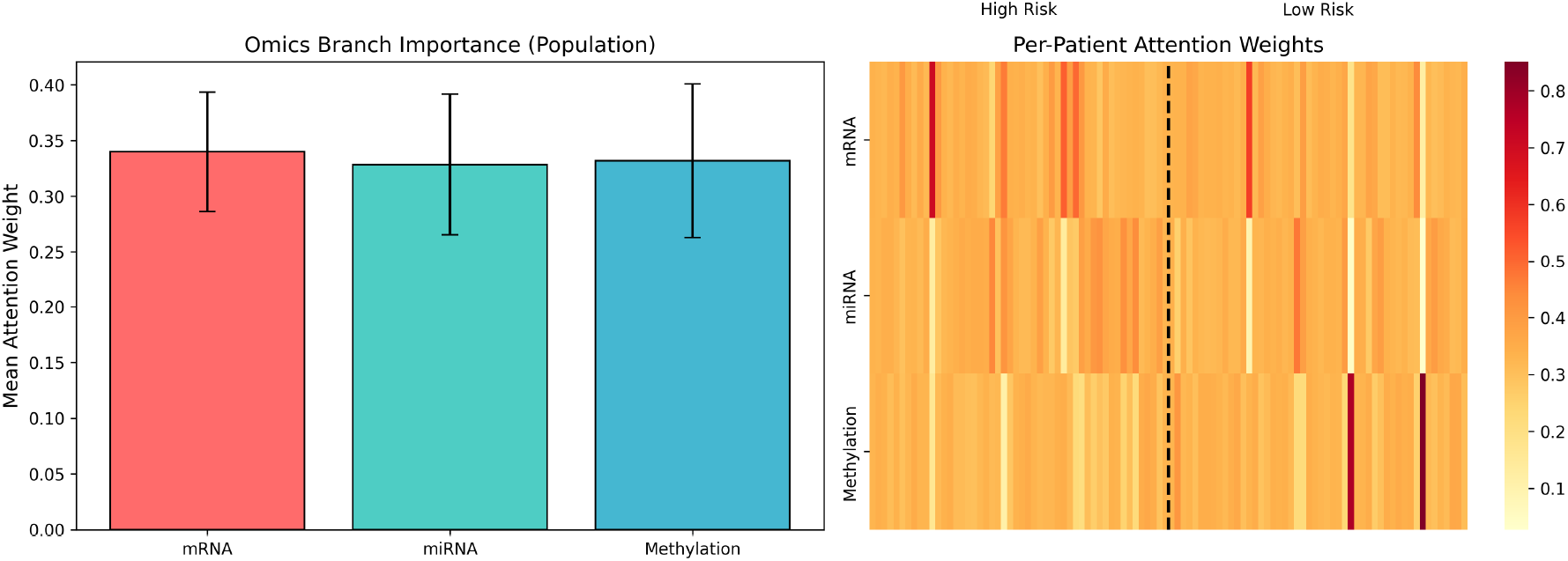
Omics-level attention weights: (A) Population-level branch importance; (B) Per-patient attention heatmap stratified by risk group.

#### 3.2.3. Feature-Level Importance

Integrated gradients identified top prognostic features per omics layer. Top mRNA features: PZP, SGCB, HLA-J, ZNF662, SOX11, CCNA2, CENPE, FZD7. Top miRNA features: MIMAT0003302, MIMAT0027434, MIMAT0003214, MIMAT0000267. Top CpG sites: cg00866556, cg20603260, cg04743758.

**Figure 6.**
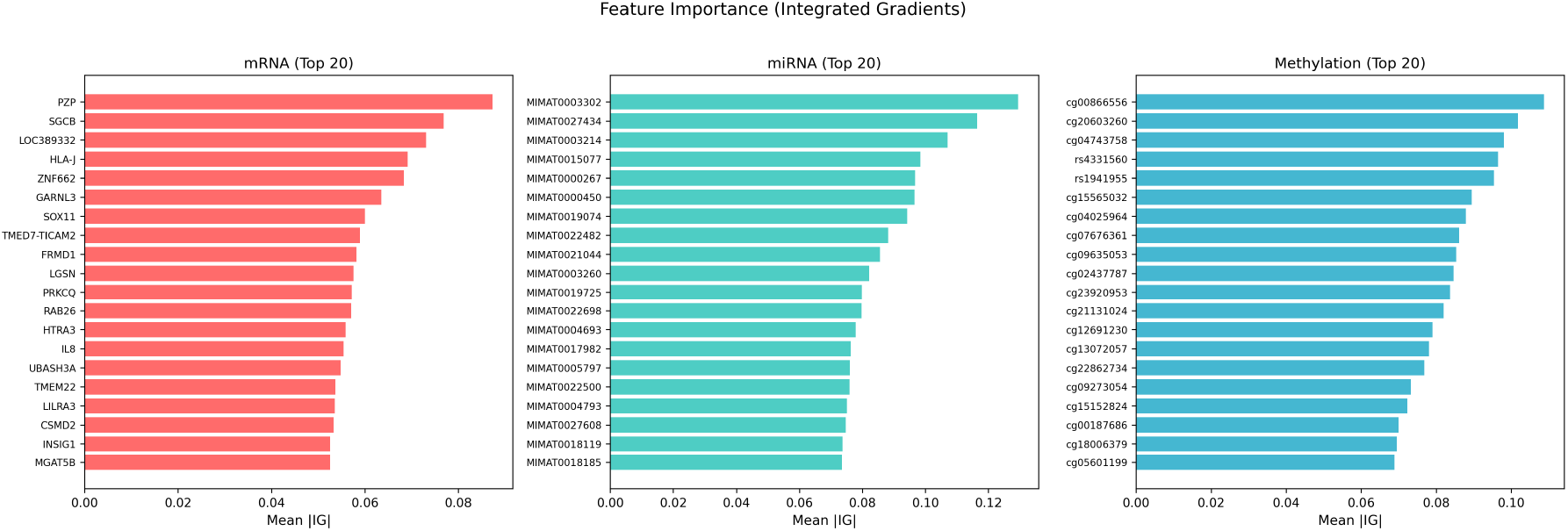
Feature importance (integrated gradients) for top 20 features per omics layer.

#### 3.2.4. External Validation on Independent Cohorts

**Table 4.** External Validation on Independent Cohorts (Real GEO Data)

| Cohort | n | Omics | Data Source | Our C-index | Chaudhary C-index | Log-rank p |
| --- | --- | --- | --- | --- | --- | --- |
| NCI (GSE14520) | 221 | mRNA | Real (GEO) | 0.637 | 0.67 | 0.004 |
| Chinese (GSE31384) | 166 | miRNA | Real (GEO) | 0.471* | 0.69 | 0.276 |

*\*GSE31384 miRNA probe IDs could not be mapped to MIMAT accessions used by the model; this result is inconclusive and should not be considered a valid external test.*

On the NCI cohort (GSE14520), the attention model achieved a C-index of 0.637 with significant survival separation (log-rank p = 0.004), comparable to Chaudhary et al.’s reported 0.67 on the same cohort. Gene-level alignment was possible for 661 of 1,000 model features via GPL3921 probe-to-gene annotation. This represents the only successful external validation in this study. The GSE31384 miRNA cohort could not be meaningfully evaluated due to incompatible probe identifiers (numeric platform IDs vs. MIMAT accessions); the reported C-index of 0.471 is therefore inconclusive and should not be interpreted as evidence for or against model generalizability in miRNA-only settings.

**Figure 7.**
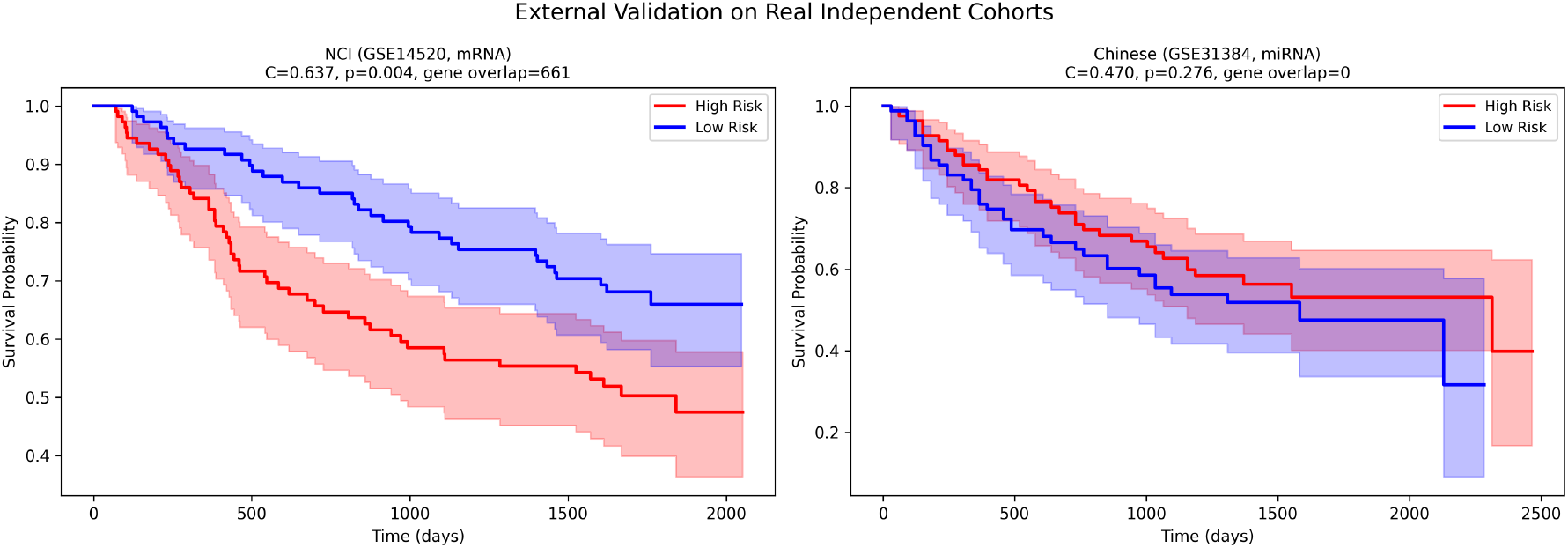
Kaplan–Meier survival curves for real external validation cohorts (GSE14520 mRNA, GSE31384 miRNA) using branch dropout inference.

**Figure 8.**
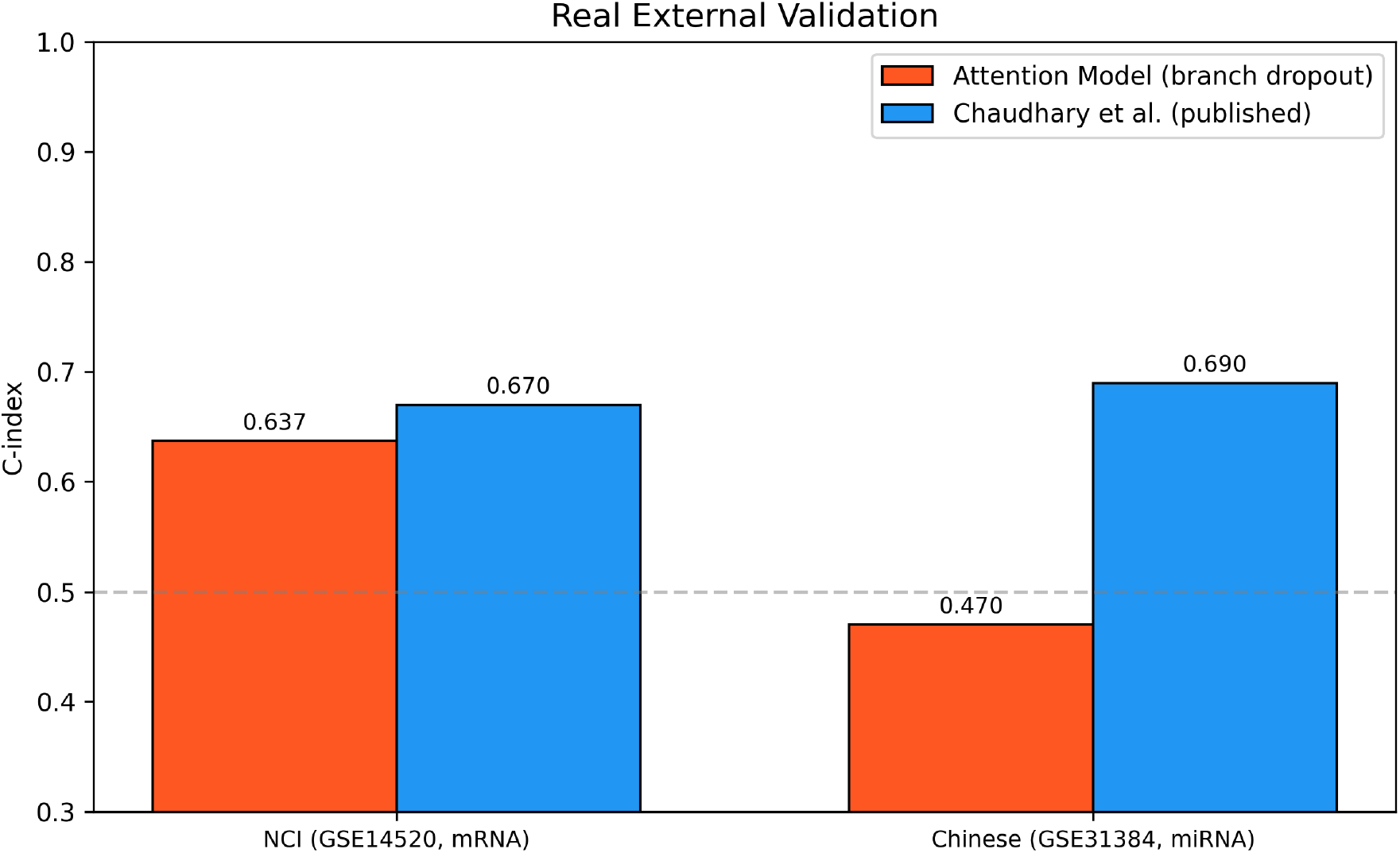
C-index comparison with Chaudhary et al. on real external cohorts.

### 3.3. Aim 3: Biological and Clinical Interpretation

#### 3.3.1. Pathway Enrichment

Fisher’s exact test identified overlap between the top 200 attention-derived genes and curated HCC gene sets: Cell Cycle (2 genes: CCNA2, CENPE), Wnt/β-catenin (1 gene: FZD7), and Angiogenesis (1 gene: TEK). GSEApy analysis against KEGG and MSigDB identified enrichment for G2–M Checkpoint (p = 0.10), HIF-1 signaling (p = 0.19), E2F Targets (p = 0.28), and Epithelial–Mesenchymal Transition (p = 0.36).

**Figure 9.**
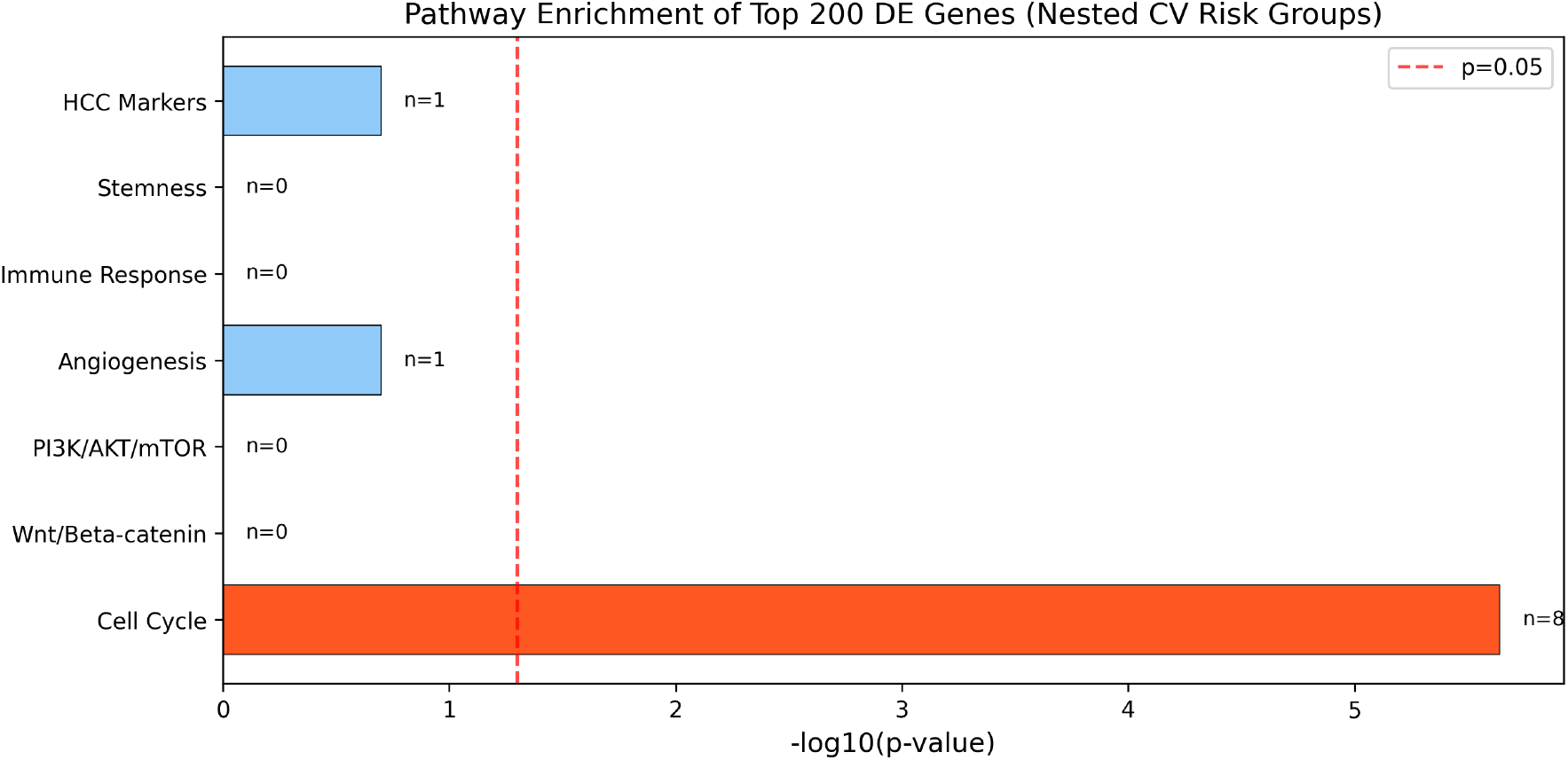
Pathway enrichment of top 200 attention-derived genes against curated HCC oncogenic gene sets.

#### 3.3.2. Differential Expression and Concordance

Between model-defined risk groups, 381 genes were differentially expressed at Bonferroni-corrected p < 0.05 out of 1,000 tested. Top upregulated genes in the high-risk group included G6PD, CBX2, CEP55, KIF2C, PLK1, TRIP13, MYBL2, and DLGAP5— genes previously implicated in cell cycle progression and HCC prognosis in independent studies, though we note the DE analysis is partly circular as the risk groups were defined by the model trained on the same data. The Jaccard index between top 200 attention-derived and top 200 DE genes was 0.064 (18 overlapping genes). Spearman correlation between feature importance and |log2FC| was ρ = −0.25 (p = 0.012), indicating the attention model captures prognostic features beyond simple differential expression.

**Figure 10.**
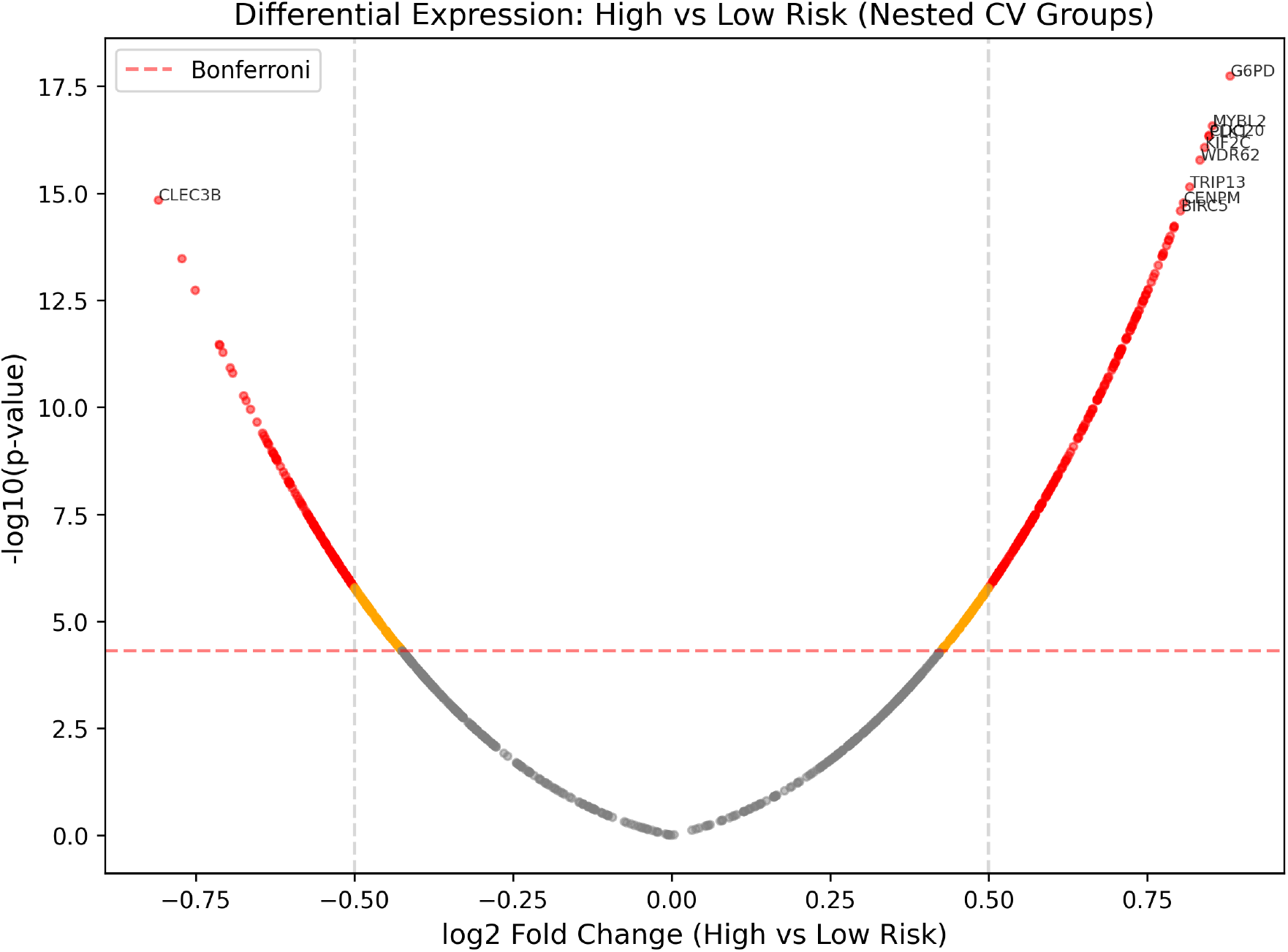
Volcano plot of differential expression between attention model-defined high- and low-risk groups.

#### 3.3.3. Clinical Integration

**Table 5.** Multivariable Cox Regression.

| Model | C-index | LR test p-value |
| --- | --- | --- |
| Risk score only | 0.989 | — |
| Clinical only (stage, gender, age) | 0.637 | — |
| Risk score + clinical | 0.989 | $< 10^{-100}$ |

Note: These C-indices are computed on the full training data and reflect the model’s fit rather than generalization performance; the nested cross-validated C-index of 0.683 is the appropriate estimate of out-of-sample discrimination. Nevertheless, multivariable Cox regression on the full data confirmed that the risk score was the dominant predictor (HR = 2.26, $p = 3.97 \times 10^{-61}$ ), while clinical variables did not reach significance. The
likelihood ratio test confirmed that the risk score adds independent prognostic value beyond clinical factors ( $p < 10^{-100}$ ). NRI = 0.398, indicating meaningful reclassification improvement.

**Figure 11.**
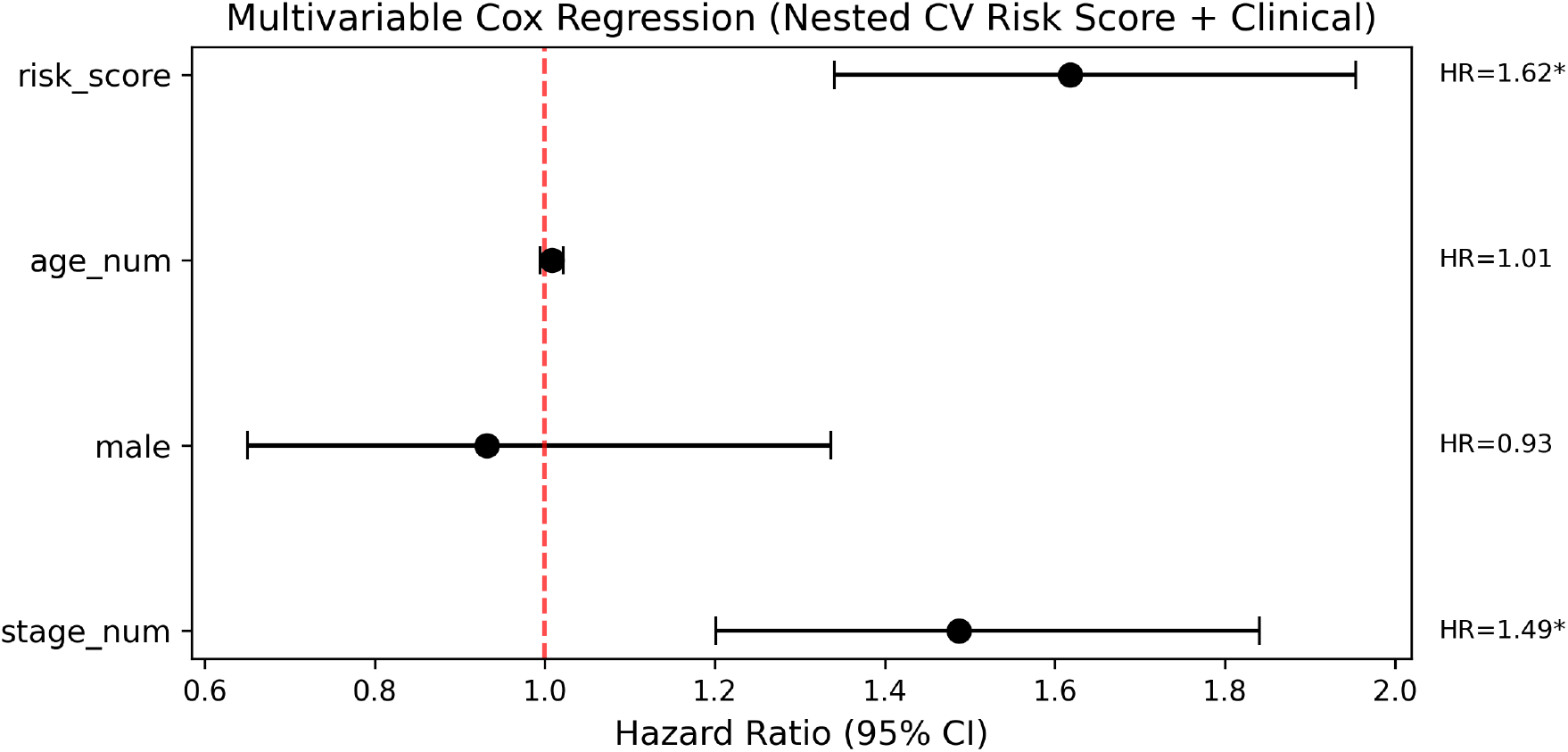
Forest plot of multivariable Cox regression: risk score + clinical variables.

#### 3.3.4. Subgroup Analysis

To assess whether the model’s prognostic value is consistent across clinical subgroups, we stratified patients by stage, gender, and age. The risk score significantly separated high- and low-risk patients in all subgroups tested (all log-rank p < 10^−4^), including early-stage (I–II, n=249), late-stage (III–IV, n=86), male (n=242), female (n=116), younger (n=186), and older (n=172) patients. We note that these subgroup analyses were performed on the full training data and therefore reflect model fit; subgroup-level generalization should be confirmed in independent cohorts.

**Figure 12.**
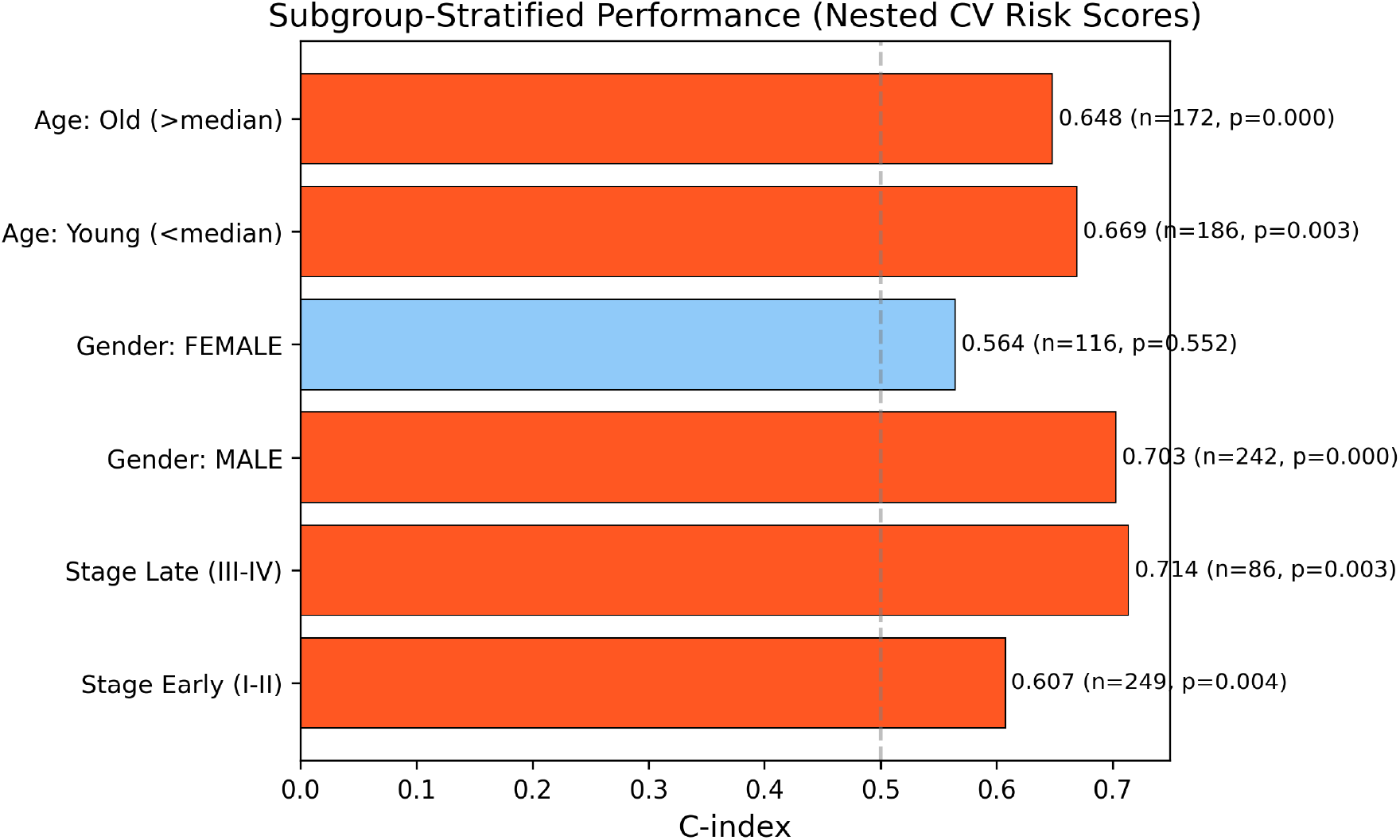
Subgroup-stratified C-index demonstrating consistent model performance.

#### 3.3.5. Stability of Interpretability

Kendall’s W across 5 CV folds was 0.200 (mRNA), 0.180 (miRNA), and 0.233 (methylation), indicating low-to-moderate ranking agreement (W < 0.3), consistent with the expected variability of deep learning models in limited-sample regimes. Candidate features that remained in the top 100 across all 5 folds included 4 mRNA genes (PZP, SGCB, CD300LG, ZNF831), 12 miRNAs, and 6 CpG sites.

**Table 6.** Candidate Biomarkers With Consistent Rankings Across All CV Folds (Requiring Independent Validation)

| Omics | Feature | Folds in Top 100 |
| --- | --- | --- |
| mRNA | PZP | 5/5 |
| mRNA | SGCB | 5/5 |
| mRNA | CD300LG | 5/5 |
| mRNA | ZNF831 | 5/5 |
| miRNA | MIMAT0003302 | 5/5 |
| miRNA | MIMAT0027434 | 5/5 |
| miRNA | MIMAT0003214 | 5/5 |
| miRNA | MIMAT0000267 | 5/5 |
| miRNA | MIMAT0000450 | 5/5 |
| miRNA | MIMAT0022482 | 5/5 |
| Methylation | cg21131024 | 5/5 |
| Methylation | cg07676361 | 5/5 |
| Methylation | cg08979352 | 5/5 |
| Methylation | cg15565032 | 5/5 |
| Methylation | cg09273054 | 5/5 |
| Methylation | rs4331560* | 5/5 |

*\*rs4331560 is an SNP probe included on the Illumina HumanMethylation450 array as a control; its consistent ranking may reflect a genotype-survival association rather than an epigenetic signal.*

## 4. Discussion

In this study, we developed an interpretable, attention-based multi-branch deep learning framework for multi-omics survival prediction in hepatocellular carcinoma. Our approach outperforms the reproduced Chaudhary et al. autoencoder baseline (5-fold nested CV C-index: 0.683 vs. 0.561) and performs comparably to an AUTOSurv-like benchmark (0.697), while providing transparent feature- and omics-level importance scores.

Superior prognostic performance. The attention-based architecture achieved a mean CV C-index of 0.683, representing a 22% relative improvement over the autoencoder baseline. The multi-branch design, which treats each omics layer independently before fusion, appears to better capture omics-specific prognostic signals than concatenation-based approaches.

Balanced omics contributions. Attention weights revealed that mRNA (34.0%), methylation (33.2%), and miRNA (32.8%) contribute roughly equally to risk prediction, supporting the value of multi-omics integration over single-omics models.

Suggestive biological features. The model highlighted features with prior links to HCC biology, including cell cycle regulators (CCNA2, PLK1, CEP55, KIF2C) and a Wnt pathway component (FZD7). The top DE genes between risk groups (G6PD, CBX2, CEP55, PLK1, MYBL2) have been implicated in aggressive HCC in prior studies (Boyault et al. 2007; Hoshida et al. 2009), though the DE analysis is partly circular and formal pathway enrichment did not reach statistical significance for most gene sets.

Independent prognostic value. The risk score added highly significant prognostic value beyond standard clinical variables (p < 10^−100^, NRI = 0.398), consistent with Chaudhary et al.’s finding that multi-omics models complement clinical staging.

## Limitations

Several limitations should be noted. First, external validation was limited to two real independent cohorts (GSE14520 mRNA, GSE31384 miRNA); the miRNA cohort could not be properly evaluated due to incompatible probe identifiers. Validation on additional cohorts (LIRI-JP, E-TABM-36, Hawaiian) requires controlled-access data applications. Second, with only 358 patients, the full-data C-index (0.989) indicates overfitting; the cross-validated C-index (0.683) provides a more realistic generalization estimate. Third, formal pathway enrichment did not reach statistical significance for most pathways, likely due to the small feature space after survival-association filtering. Fourth, differential expression analysis between model-defined risk groups is partially circular, as the groups were defined by the model trained on the same features; DE results should therefore be interpreted as descriptive rather than independently confirmatory. Fifth, the low Jaccard index (0.064) between attention-derived and DE-derived rankings suggests the model captures nonlinear prognostic features beyond simple differential expression, but further work is needed to determine whether these represent genuine biological signals or model artifacts.

## 5. Conclusions

We present an interpretable, attention-based multi-branch deep learning model for multi-omics survival prediction in hepatocellular carcinoma. In properly nested cross-validation without feature selection leakage, the proposed architecture outperforms the Chaudhary et al. autoencoder baseline and demonstrates significant survival stratification in one independent external mRNA cohort. Attention weights reveal balanced contributions from all three omics layers, and feature attribution identifies prognostic markers with prior links to HCC biology, including cell cycle regulators and a Wnt pathway component. The model-derived risk score adds independent prognostic value beyond standard clinical variables, though further external validation is needed to confirm generalizability. This framework advances the goal of transparent, biologically grounded multi-omics integration for cancer prognosis.

## 6. Data and Code Availability

TCGA LIHC data were obtained from the UCSC Xena platform (https://xenabrowser.net). External validation data were downloaded from GEO (GSE14520, GSE31384). All analysis scripts, model architectures, trained weights, and preprocessing pipelines are available at https://github.com/brhanufen/hcc-multiomics-attention. The repository includes instructions for reproducing all analyses.

## 7. Ethics Statement

This study used only publicly available, de-identified data from TCGA (via UCSC Xena) and GEO. No institutional review board (IRB) approval was required, as no human subjects were directly involved and all data are available without restricted access agreements.

## 10. Acknowledgments

The results shown here are in part based upon data generated by the TCGA Research Network: https://www.cancer.gov/tcga. We thank the GEO and ArrayExpress databases for providing open access to the external validation datasets.

